# A knowledge graph and topological data analysis framework to disentangle the tomato-multi pathogens complex gene regulatory network

**DOI:** 10.1101/2025.04.09.647963

**Authors:** Maxime Multari, Mathieu Carrière, Xavier Amorós-Gabarrón, Alexina Damy, Sebastian Lobentanzer, Julio Saez-Rodriguez, Stéphanie Jaubert, Aurélien Dugourd, Silvia Bottini

## Abstract

Global population is rapidly increasing, representing a major challenge for food supply, exacerbated by climate change and environmental degradation. Despite the pivotal role of agriculture, plant health and survival are threatened by various biotic stressors. Although how plants respond to each of these individual stresses is well studied, little is known about how they respond to a combination of many of these bio-aggressors occurring together.

To tackle this question, first, we built TomTom, a knowledge graph gathering molecular interactions from nine publicly available databases, including transcription factors- or microRNAs-targets, protein-protein interactions, and functional terms. Then, we selected transcriptomics data of tomato subjected to six distinct pathogens and performed an integrative analysis. We found 5561 candidate genes involved in the multi-stress response of tomato. To study how the response is orchestrated, we mapped those genes in TomTom and extracted a comprehensive gene regulatory network (GRN) composed of 71 transcription factors (TF) and 1786 target genes. By estimating the TF activity, we identified 43 TFs responding either specifically to one or multiple bio-aggressors. GRN analyses with a topological data analysis approach allowed to identify 18 clusters of TFs with similar properties, yielding four main configurations localized in specific regions of the GRN. Finally, we found one NAC and four ERF hubs which cooperatively coordinate the tomato response to multiple pathogens.

Our findings allowed to study the complex molecular reprogramming in tomato upon interaction with different biotic agents, providing tools scalable to other questions involving tomato molecular interactions and beyond.

## Introduction

Plants as sessile organisms are exposed to diverse stresses, either abiotic or biotic. These stresses can act singly, in combination, sequentially or in a multifactorial manner [1], and their combination can have dramatic consequences for plant survival even when the effect of each stress applied individually is negligible [2] [3]. Plant pathogens have a broad spectrum of characteristics spanning from ectoparasites to endoparasites, from biotrophic to necrotrophic, from avirulent to virulent strains, attacking different tissues from leaves to roots, varying time-life cycles, causing several different possible diseases that can lead to different plant responses [4]. Although multiple infections affecting a single plant or crop are recognized to be common in plant disease epidemics [5], plant diseases are usually studied as the result of the interaction between a plant host and a single pathogen. Single stress studies in model systems have provided invaluable mechanistic insights, but more sophisticated experimental designs are required to replicate the native context of a plant. However, such studies can be expected to be more difficult to perform and interpret.

Advances in systems biology have facilitated the understanding of several complex biological processes including plant immune systems, notably using graph-based techniques. Such techniques rely on an abstract description of molecular interactions as nodes’ relationships (edges) [6] where nodes can represent molecules such as genes, RNAs, proteins, metabolites, etc. [7]. Network inference has emerged as a broadly used methodology with the development of the so-called omics techniques and their spectacular progresses in the last decade [8]. Among those, transcriptomics is by far the most produced omics type, offering the possibility to study transcriptional regulation, an important factor in controlling tolerance and resistance to stresses. Therefore, Gene Regulatory Networks (GRNs) composed of nodes which represent transcription factors (TFs) or target genes, and edges representing the regulatory connections between regulators (TFs) and targets (genes), have gained a lot of attention lately [9]. GRN inference is an evergreen unsolved problem in the field of systems biology. Two main approaches are commonly employed, either using omics data to perform a data-driven inference [10] or using prior knowledge, such as known molecular interactions from publicly available databases [11], to build a network skeleton. While reconstructing interactions from observational data is a critical need for investigating natural biological networks, the dimensionality is usually very high. On the other side, summarizing knowledge of an organism is challenging, most of the time relevant information is found across multiple databases, often not interoperable among each other.

To tackle these challenges, we focus on tomato (*Solanum lycopersicum*) not only because of its agroeconomic interest, but mainly because it is susceptible to more than 200 diseases caused by several different pathogens [12], [13] [14] [15], offering a biological model to study multi stress integration. First, we developed TomTom, a knowledge graph for molecular interactions in tomato. Then, we retrieved publicly available transcriptomics data of tomato subjected to six different pathogens. We extracted the molecular signatures which are either specifically responding to one stress or to multiple ones by performing an integrative analysis with the HIVE tool [16]. We used TomTom to generate the GRN including only the molecular signatures extracted from our integrative approach. To study this complex network, we set up a novel approach based on the estimation of transcription factor activity [17] and the study of the topology of the network using the topological data analysis framework [18].

## Materials and Methods

### TomTom: a knowledge graph to study tomato molecular interactions

Several databases were used to construct the knowledge graph to cover multiple molecular interactions. Supplementary Table 1 summarizes all the input databases. The procedures used to include them in the knowledge graph are explained in the following paragraphs.

### Genome and microRNA backbone

The genome of *Solanum lycopersicum* was downloaded from Sol Genomics Network [19], using the latest version available: SL4.1. The GFF file, describing genomic features, was used as a backbone for the knowledge graph to construct and link the genomics entities (genes - transcripts - proteins), using unique identification. The identification (ID) chosen was the Organized Locus Name (OLN) of each gene (eg: Solyc01g010203). This ID is constructed thanks to the species name (*SOlanum LYCopersicum*) followed by the chromosome number (01). A letter (g) specifies the type of the entity, here a gene, followed by a unique 6-number identifier (010203). We extracted the genes from the GFF file and followed the central dogma of biology to add transcripts and proteins. Thus, for one gene we have one transcript and one protein, as alternative splicing was absent in the reference genome. The previous features build the backbone of the knowledge graph by linking genes to transcripts and transcripts to proteins.

Then, we also constructed the microRNA backbone (microRNAs and precursors) using miRbase [20].

We downloaded miRBase and parsed it to retrieve the microRNA and precursor information for *S. lycopersicum* species. We then linked each precursor to its corresponding microRNA.

### Integration rules for other databases

We had to define specific rules to integrate other databases that cover multiple interactions.

Since all these databases are based on different genome versions, to have consistent interactions when considering different genome versions, a filtering process was mandatory to avoid adding incorrect information to the graph. Thus, we checked every added entity against the backbone to avoid having ones not present in the genome used (SL4.1). All identifiers present in the input databases must match the identifiers in the backbone, and if one of the identifiers is not recognized in the backbone, the corresponding interaction is removed.

The filtering rules have been applied to all further databases.

### Transcription Factor - Target Interaction

The transcription factors (TFs) and their associated family were downloaded from the PlantTFDB [21] web interface and added to the graph. To connect TFs and target genes, we downloaded the gene regulatory network (GRN) of *S. lycopersicum* from the PlantRegMap [22] web interface. The GRN, composed of TF-Target interaction, was added to the graph.

### MicroRNA - Target Interaction

Three microRNA - Target interaction (MTI) databases: TarDB [23], DPMIND [24], and PNRD [25] were added by downloading the interactions from their respective web interfaces.

PNRD also included precursors and not only mature microRNA IDs. We manually curated those cases by checking whether they were present in miRBase using their sequence and renamed those microRNAs accordingly or filtered them otherwise.

The three databases are distinctly represented in the graph, giving database-specific information and the ability to perform specific queries that include one or more MTI databases.

### Protein - Protein Interaction

Protein-protein interactions (PPI) were retrieved from the STRING database [26]. STRING uses its identifier, derived from UniProt, instead of the OLN identifier. To associate an OLN, we used the UniProt mapping script that maps the protein’s IDs to the associated reference genome. However, the only available version of the *Solanum lycopersicum* genome is the SL3.1 from Ensembl Plants. The drawback of this approach is that by mapping on a different genome version than the one used for the KG, there is a loss of proteins. After converting the protein IDs to the OLN format, the PPI network was added to the knowledge graph. We included interactions with a STRING combined score greater than 0.8 to maintain high-quality interactions. The combined score can be adjusted by the user if more PPIs are to be added.

### Planteome Terms

To gain more insights into the gene functions, we downloaded annotation data from Planteome [27], which included terms from Gene Ontology (GO) or Plant Ontology (PO) as well as the term descriptions. Those terms were then linked to genes, accordingly, using a custom procedure to match genes to terms by reformatting the original downloaded file.

### KEGG Pathways

KEGG pathways for *S. lycopersicum* were retrieved through KEGG API [28]. Precisely three requests: pathway IDs and names, genes involved in pathways, and mapping from KEGG gene IDs to UniProt IDs. UniProt IDs were then converted to OLN using the same script as before, provided by UniProt. Once converted, links between genes and KEGG pathways were created.

### OMA orthologs

OMA orthologs of *Arabidopsis thaliana* for the *S. lycopersicum* species were retrieved using the OMA REST API python package [29]. The *A. thaliana* orthologs were linked to the genes according to the database information.

### Knowledge graph construction

To construct the knowledge graph, we used the BioCypher (v0.5.40) framework [30]. This allowed us to systematize Python modules for integrating different files or databases into the graph and add complementary information, including node IDs, node properties, edge IDs, and edge properties. The semantics used in the graph were defined in a YAML file that maps graph entities and relationships to concrete concepts from an ontology.

For each database, a corresponding Python module (“adapter”) was created to consider the peculiarities of each database in terms of the information and type of interactions.

The graph was then constructed by running a pipeline orchestration script that controlled each step of the construction. The knowledge graph was finally deployed into a Neo4j graph database instance via a Docker Compose workflow, created from BioCypher’s template.

### Integrated multi-transcriptomics dataset composition and HIVE analysis

The integrated data comes from five different publicly available studies and is described in Supplementary Table 2. For each project, we selected the corresponding sequencing data from the tomato interaction with pathogens or controls. A summary of all the samples used is described in Supplementary Table 3. In total, 83 samples were included in the dataset, representing seven *S*.*lycopersicum*-pathogen interactions. Each sample was preprocessed using an internal pipeline using MultiQC [31] then trimmed using fastp [32] with parameters: -5 -3 -M 28 -l α -y. The α is set to be the read length - 2 for each sample. The sequences were then mapped to the *S. lycopersicum* v.4.1 genome using STAR [33] with the following parameters: for the indexing: --runMode genomeGenerate – genomeSAindexNbases 10; and the mapping: --alignEndsType EndToEnd – twopassMode Basic. We further counted the reads on the alignment using featureCounts [34] with default parameters. We obtained a count matrix for each dataset. We merged all the count matrices to obtain one multi-transcriptomics dataset.

We further applied HIVE [16] to the multi-transcriptomics dataset with default parameters except for the number of bins, which was set to 138 in the “shap_bin_selection_f()” function. HIVE outputted a list of 5561 genes.

### Wald statistic calculation

Differential expression analysis was conducted using DESeq2 [35] with the default parameters. Each dataset was analyzed independently, and we compared the controls (uninfected) to the infected samples. The Wald statistics, calculated by dividing the log2FC by its error (lfcSE) for each gene, are used for further analysis.

### Gene Regulatory Network inference

To retrieve the GRN from the HIVE results list, we queried TomTom to have both TF and targeted genes coming from the list. We removed the low-confidence “motif” evidence-only interactions.

### Transcription factor and pathway activity estimation

The transcription factor (TF) activity was computed using the Univariate Linear Model (ULM) from the decoupleR [17] python package on the GRN.

For each sample, ULM fits a linear model for each TF with the stats of targeted genes being the response variable and the TF interaction classes (activating or repressing TF) being the explanatory variable. The t-value and the p-value of the fitted model are computed. The predicted activity of the TF is based on the t-value, representing an active TF (positive value) or an inactive one (negative value), also referred to as TF activity score. In other words, the predicted activity reflects if a TF follows the expression pattern of its targets (active TF) or not (inactive TF).

Since we did not have information on TF classes, all TFs in the GRN are considered activators. Therefore, the activity is dependent on this activating class assumption for all the regulator TFs in the GRN. By applying this model, we retrieved TFs with a p-value inferior to 0.05 in at least one condition. The pathway activity was computed using the Multivariate Linear Model (MLM) from decoupleR [17]. The network used in this case links genes from the GRN to KEGG pathways, as previously added in the KG. As for the GRN, we missed the interaction classes between genes and pathways; thus, every interaction is considered positive. Like before, the stats of each gene in pathways are used to compute the activities. We then retrieved pathways with a p-value inferior to 0.05 in at least one condition.

### Topological data analysis

In order to study large networks of TFs, we use the Mapper algorithm [36] from topological data analysis in combination with the mode-based clustering algorithm ToMATo [37].

### Mapper algorithm

This algorithm is a data visualization method that produces a graph representing the dataset, and whose topology (connected components, branches, loops) matches that of the original data. It can be obtained from a continuous function, called filter *f*: *X* → *R*^*d*^, where *X* is the dataset. Let *I* be a cover of the image of the filter *im*(*f*) with overlapping hypercubes: *im*(*f*) ⊆ U_*H*∈*I*_ *H*. This cover can be pulled back to *X* using *f*^−1^; this induces the pullback cover *U* {*f*^−1^ (*H*) *X* | *H* ∈ *I*}. The pullback cover is then refined by separating every cover element of into its connected components with a clustering algorithm, this is called the connected pullback cover:

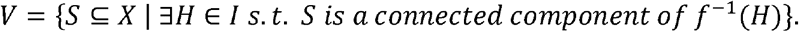

Finally, the Mapper graph is defined as the nerve of *V*, that is, the graph whose nodes are in bijection with the elements of *V* (each Mapper node thus represents a group of similar data points), and whose edges are drawn between two nodes as soon as the corresponding cover elements in *V* have at least one data point in common (Supplementary figure 1A).

We applied the Mapper algorithm on the GRN using the first two eigenvectors of the GRN normalized Laplacian as filter (i.e., the two eigenvectors with largest eigenvalues), and a cover of 25 hypercubes with 30% overlap, evenly sampled along the filter components. The clustering algorithm used for producing the connected pullback cover was defined as the connected components of the GRN restricted to the preimages of the hypercubes under the filter. Finally the Mapper nodes were colored using the average activities of the significant TFs in response to each pathogen, and the Mapper edges were thickened according to the number of target genes shared by the TFs in the corresponding Mapper nodes. We also removed 2 nodes (10 and 12) from the final graph because they were isolated and without any significant TFs. This allowed us to produce compact and geometric representations of the GRN.

### ToMATo clustering

In order to group TFs together, we applied the Topological Mode Analysis Tool (ToMATo) clustering algorithm on the Mapper graph. ToMATo is a hierarchical clustering algorithm which uses a user-defined scalar-valued filter *f*: *X* → *R* in order to build the cluster hierarchy. Specifically, this algorithm first requires building a graph *G* on top of the dataset (e.g., a neighborhood graph), and then uses this graph to approximate the gradient *∇f*(*v*) at every graph node *v* ∈ *G*. Finally, the clusters are computed by capturing all local maxima of *f* using gradient flows (each maximum being seen as a new cluster), and a hierarchy is built by looking at the saddle points of *f* and connecting all the clusters associated with the saddle to the one with highest prominence. This method allows us to define hierarchies of clusters for any arbitrary function and is known to be robust to noise in the data.

Hence, we applied the ToMATo clustering algorithm using the Mapper graph defined above as the graph *G*, and the TF activities as well as edge thickness as filters, after some pre-processing has been applied, leading to 8 different clusterings. The pre-processing used was obtained by applying the following mixtures of Gaussians to the TF activities and the edge thickness, showed in Supplementary figure 1B.

Indeed, this pre-processing on the TF activities (resp. edge thickness) allowed us to characterize TFs with low, high, and null activities in average (resp. TFs with no common target genes and many common target genes) as local maxima of the new filter. Finally, the eight clusterings were combined together using Cartesian products, and simplified in the following way: if the TF activities of a Mapper node are significant in response to at least two different pathogens in the Cartesian-product cluster corresponding to the Mapper node, then this node is marked as “multiple response”, otherwise it is marked as “specific response”, and if the edge thickness cluster of the Mapper node is associated to a maxima that is below the saddle point of the Gaussian mixture, it is marked as “sparsely connected”, otherwise it is marked as “highly connected”. We further identify groups of nodes having the same label, allowing us to create four subparts of the original graph.

### Hub identification

We performed hub identification in each group found by the ToMATo algorithm by retrieving the sub-network composed of the TF and targets of these groups and computing two metrics: degree and betweenness. We defined a hub as a node that have the maximum values for one or both metrics. If multiple nodes have the same value, we considered all of them to be hubs. We retrieved only the hubs with significant TF activity.

## Results

### TomTom: a comprehensive knowledge graph summarizing molecular interactions in tomato

To build the knowledge graph TomTom, we collected different types of molecular interactions and annotations from eleven databases (Supplementary Table 1). The construction of the graph relied on a backbone of 34688 unique genes present in the SL4.1 tomato genome. Interactions were then added following the gene-transcript-protein schema (Figure 1A). For the microRNA backbone, from miRbase [20] 112 precursors and 147 microRNAs were included in the graph. The microRNA-target interactions, were retrieved from DPMIND [24], TarDB [23] and PNRD [25] (318, 1554 and 616 connections, respectively). From the genes, 1717 transcription factors (TFs) were identified from PlantTFDB [21] and by using PlantRegMap [22], we included 482776 TF-gene interactions in the TomTom. Then, 226944 protein-protein interactions were recovered from STRING [26]. Annotation terms were selected from Planteome [27], (6551 terms), consisting of 267480 links between genes and terms, and 159 pathways and 14916 links from KEGG [28]. From OMA [29] we retrieved 13342 orthologs in *Arabidopsis thaliana* and included 35830 relationships in TomTom. In total, TomTom encompassed 124375 nodes and 1169480 relationships (Figure 1B and C). The automatic pipeline can be rerun at any time to provide an updated graph. The graph can be queried using the Neo4j graph database query language, Cypher.

**Figure 1.**
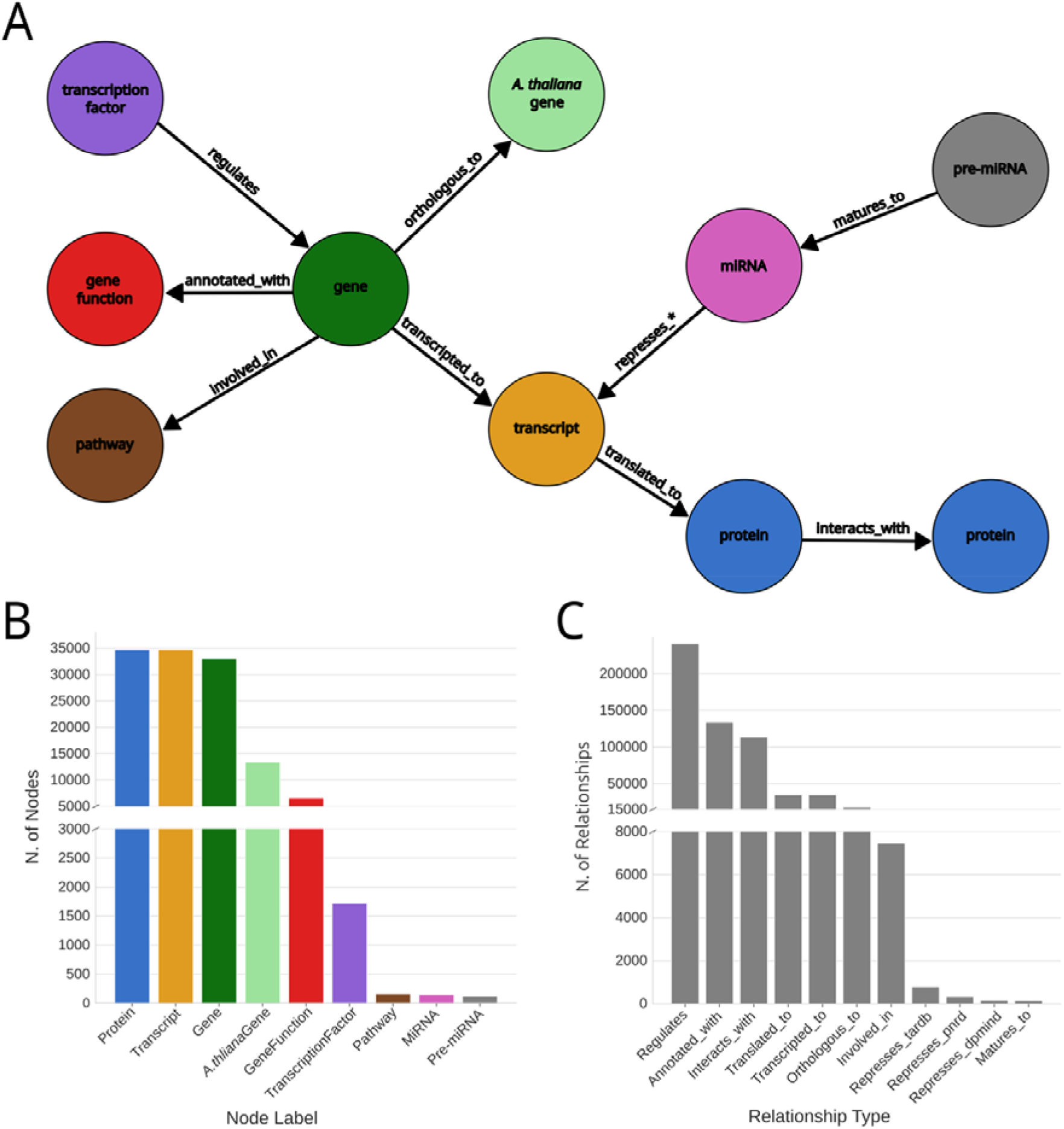
TomTom schema. A) Type of node entities and their relationships in TomTom. Each node represents an entity, and the nodes relate to associated relationships. The * in the “represses_*” interaction represents the database name, such as pnrd, dpmind, and tardb. B) Number of nodes by molecule type or function. C) Number of edges by relationship type.

Overall, TomTom provides a global summary of *Solanum lycopersicum* from molecular interactions to pathway regulation.

### The multi-stress transcriptomic data integration and TomTom allowed to infer a global GRN, modeling tomato response to single and multiple attacks

To study the tomato response to multiple biotic stresses, we collected transcriptomics data from publicly available studies, each challenging tomato plants with one distinct pathogen, infecting either the leaf or root (Supplementary Table 2). The leaf pathogens include the fungi *Botrytis cinerea* and *Cladosporium fulvum*, the oomycete *Phytophthora infestans*. The root pathogens include the root knot nematode *Meloidogyne incognita* (two time points: 7 and 14 days post infection, dpi) and a mild and a severe strain of potato spindle tuber viroid (PSTVd). The choice of those pathogens was made to include the highest variety of species, lifestyle and tissue attacked. The integrated analysis of the data was performed using HIVE [16].

To study how regulatory processes are modulated specifically or to respond to multiple pathogens, we built a gene regulatory network (GRN). Briefly, we queried TomTom to extract all relationships corresponding to “TF-target”, where targets can be either genes or TFs, involving only genes found in the HIVE results, and we excluded the interactions supported only by motif screening analysis. The final GRN consists of 1786 nodes including 71 TFs and 4201 edges (Figure 2A, Supplementary Figure 2). First, we performed a KEGG pathways activity analysis of the GRN by using a multivariate linear model on the t-values of the genes for each condition belonging to the examined pathway (see methods) [38]. Among the significantly active pathways, we found: plant-pathogen interaction, oxidative phosphorylation, and metabolism related terms. By looking at the overall pattern of pathway activities, we can observe that the more similar the characteristics of the bio-aggressors, the more similar is the tomato response to the pathogen attack (Figure 2B). The response to *M. incognita* is apart and looks overall opposite to the response to the other pathogens, which correspond to the very different mode of infection of this pathogen compared to the others included in this study (Supplementary Fig 3). We observe that six out of 13 pathways are active in response to a specific pathogen and none of them is active in response to all pathogens (Figure 2C). From Figure 2D, we observe that leaf pathogens trigger predominantly negative activated pathways, while root pathogens yield a majority of positive ones. To better quantify the number of pathways activated in response to multiple pathogens and their activity levels and significances, we represented those results using a network representation (Figure 2E).

**Figure 2.**
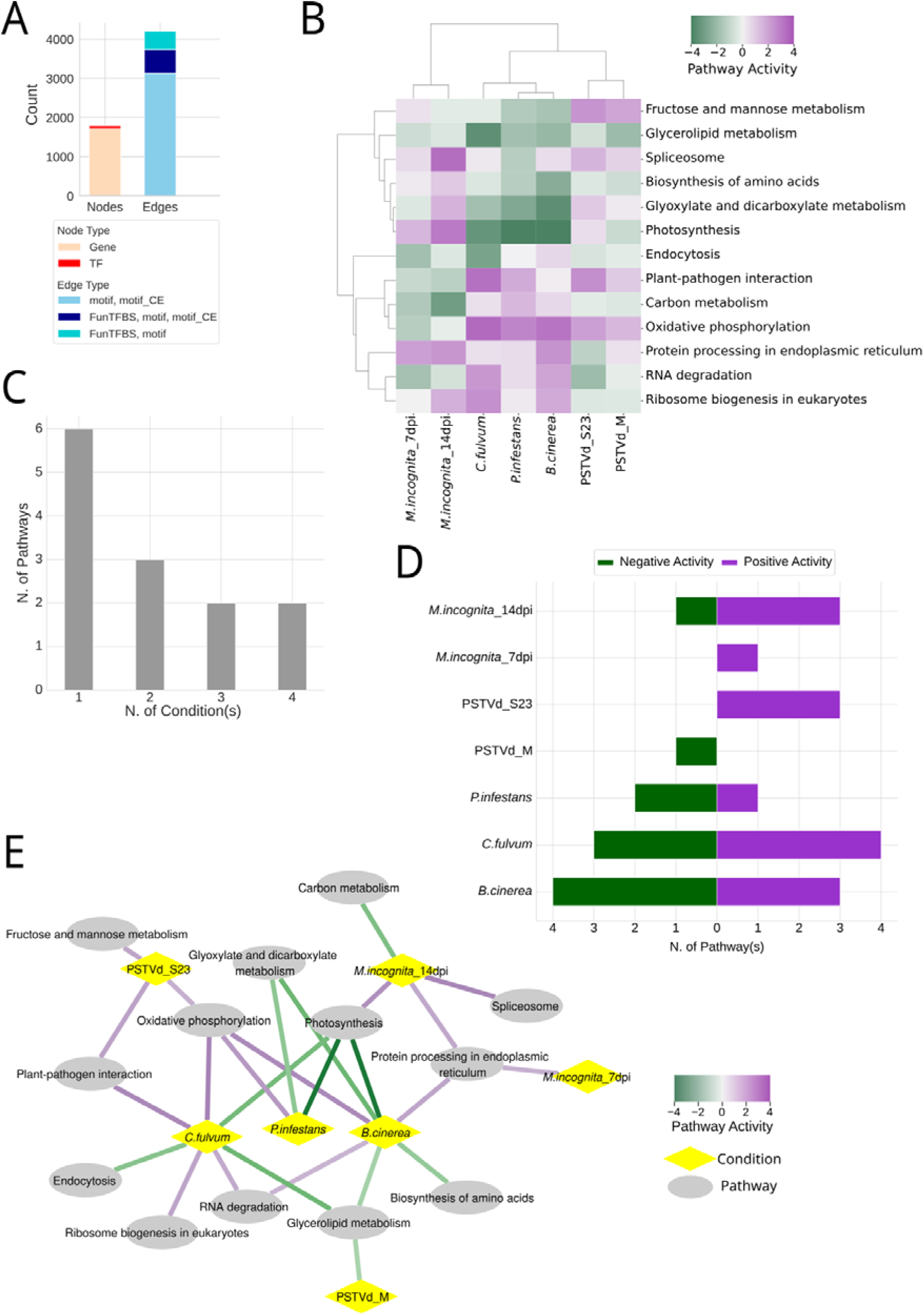
Characteristics of the GRN inferred by TomTom and the transcriptomic integrative analysis of tomato subjected to multi-stress. A) Number of nodes or edges by type in the GRN. B) Clustermap showing the levels of pathway activity in each condition only for significant results in at least one condition. C) Barplot reporting the number of pathways with significant activity in one or more combination of conditions. D) Barplot showing the number of pathways with significant positive (on the right) or negative (on the left) activity according to conditions. E) Network representation of the pathway activity analysis. Nodes are either pathways with significant activity or conditions, edges represent significant activity in the connected condition, the activity value is reported as color scale of the edges.

Overall, we observe that the three leaf pathogens trigger the activation of shared pathways in tomato, suggesting that there is a core common response to those pathogens. We also found pathways activated in response to pathogens with different characteristics, although with different sign of activity. This supports the hypothesis that different pathogens can trigger the same pathways but with different modulation.

### Estimating the transcription factor activity identified 43 TFs orchestrating specific and multi-factorial tomato response to biotic stressors

TFs included in the GRN belong to 23 different families. Among the families with the highest number of TFs in the GRN we found bZIP and ERF (8 TFs each), MYB and NAC (6 TFs each), C2H2 with 5 TFs and WRKY with 4 TFs (Figure 3A). Since the expression of TFs is not always linked to their regulatory activity, we estimate the putative activity of all TFs in each condition based on the expression of their regulated targets, similarly to the pathway activity estimation explained in the previous paragraph (see methods) [17]. This analysis yields 43 TFs with a significant activity in response to at least one pathogen infection. As shown in Supplementary Figure 4A, the TF activity is not correlated to the t-value (correlation range: -0.18 to 0.21 and we found that there is no relationship between the number of targets and the activity of TF, therefore there is no detectable size effect in the TF activity estimation (Supplementary Figure 4B).

**Figure 3.**
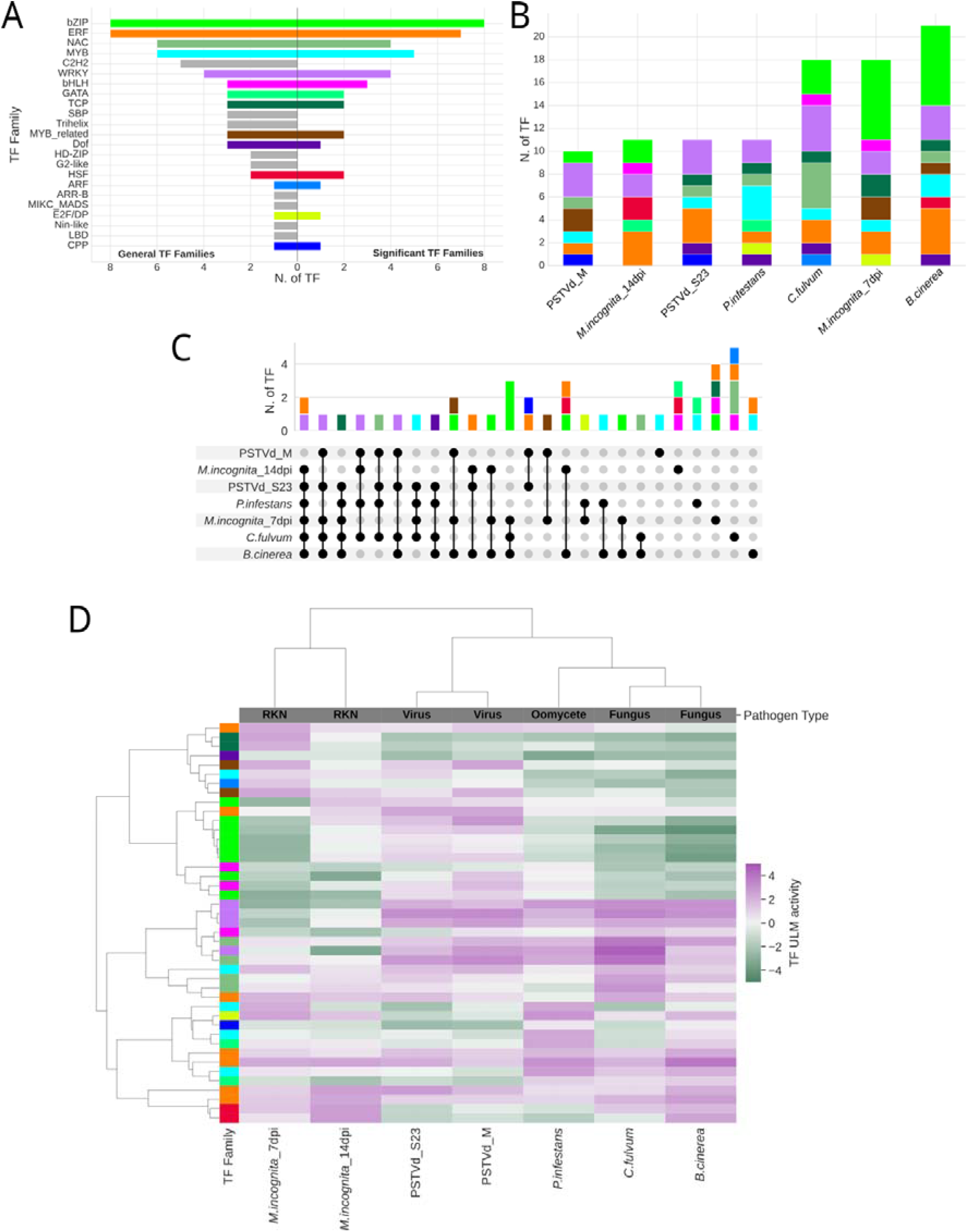
TF activity. A) Number of TF in the GRN per family (on the left) and only for significant activity (on the right). B) Number of TF with significant activity by family, accordingly to A, per condition. C) Upset plot showing the number of conditions in which TFs were reported to have a significant activity. Colors accordingly to TF family, legend in A. D) Heatmap and hierarchical clustering, showing the pattern of TF activity (only for significant in at least one condition) in response to each pathogen infection. The family of the TF is reported on the left, colored accordingly to A.

The TFs with a significant activity belong to 14 families, ruling out 9 families present in the GRN as they do not play a significant regulatory role in response to the studied biotic stresses (Figure 3A). We then focused on studying the relationship between the family and the condition in which TFs are significantly active (Figure 3B). Infection of either *C. fulvum, B. cinerea* and *M. incognita* (7 dpi) activates the highest number of TFs and two families responding only to one specific pathogen attack, i.e. CPP to viruses and ARF to *C. fulvum* (Figure 3B). We found that 17 TFs were reported active to respond specifically to only one stressor and nine are active in at least four conditions (Figure 3C). The four WRKY TFs are active in at least 4 conditions, suggesting that TFs belonging to this family might have a potential broad role in regulating transcription activity in response to multiple pathogens (Figure 3C). Opposite scenario for the bHLH TFs, which are active in response to specific pathogens. The overall activity profiles of TFs in the different conditions led to similar results as observed for the pathway activity (Figure 3D, Supplementary Figure 5).

In conclusion, the GRN inferred by combining TomTom knowledge graph and HIVE selection allowed to estimate TFs activity in response to specific or multiple biotic stressors.

### The topological data analysis of the GRN yielded four configurations linking the network structure and the TFs function

After exploring the global GRN signatures, we decided to focus on the analysis of the GRN for each pathogen response. To extract those patterns, using the mapper algorithm [36] from topological data analysis (TDA) [18] (see methods), we obtained a simplified and compressed version of the original GRN, composed of 18 nodes, each node represents a cluster of TFs (Figure 4A, Supplementary Figure 6.). The clusters in the upper and central regions are active in response to all attacks, although with different activity, while the clusters in the bottom part are active in response to specific pests. Infections by the three leaf pathogens lead to the activation of several clusters, compared to the other root pathogens studied. *C. fulvum* or *B. cinerea* trigger similar TFs responses with predominantly negative activity in the clusters in the upper region, and positive in the bottom right. The viruses seem to trigger an opposite response with a mainly positive activity in the upper part of the network. Clusters with TFs involved in *M. incognita* response have mainly a negative activity, especially in the upper region where are most of TFs involved in this response. Finally, the cluster at the bottom left is negatively active only in response to this pathogen and *B. cinerea*.

**Figure 4.**
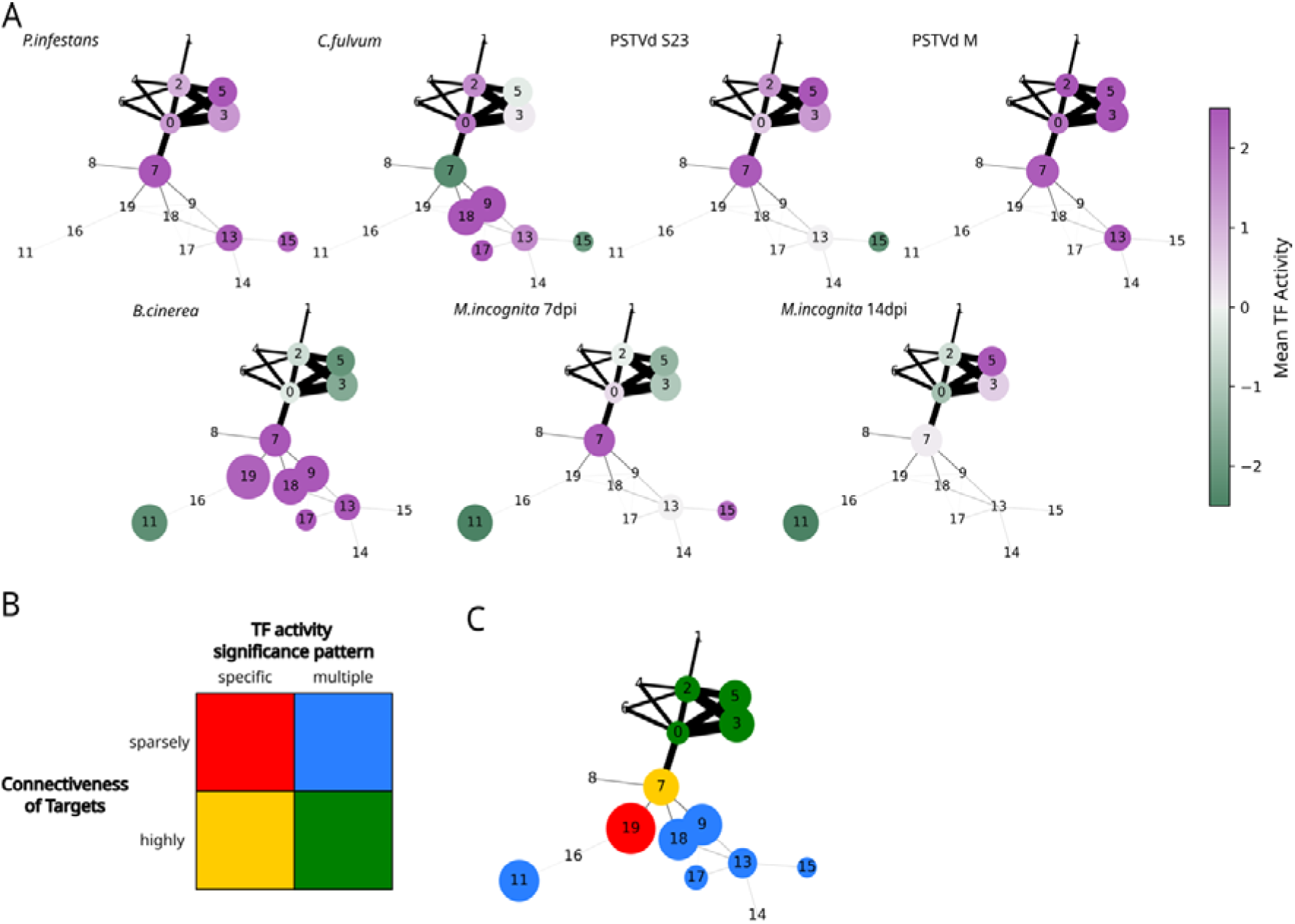
The Mapper algorithm from TDA yielded a simplified and compressed version of the GRN. A) The novel GRN representation composed of 18 nodes, each node represents a cluster of TFs, where the node size represents the number of targets in each cluster and edges represent shared targets among clusters and the edge size is proportional to the number of shared targets, nodes are colored by average TFs activity in clusters per pathogen infection. B) The four configurations representing different combinations of network structure and TFs activity. C) The four groups of clusters found by the Mapper and ToMATo algorithms from TDA without *a priori* which correspond to the four configurations in B.

The novel representation of the GRN obtained with the Mapper algorithm highlights that there are two different structures: clusters with a high number of targets but few connections with close clusters and clusters with similar number of intra/extra connections. Regarding the activity, we have two main behaviors, either TFs respond specifically to only one attack or to multiple stresses. Therefore, we envisage four different possible scenarios: “sparsely connected targets, specific response”, one or a group of TFs with specific targets, responding to a specific attack; “sparsely connected targets, multiple response”, one or a group of TFs with poorly connected targets, responding to different attacks; “highly connected targets, specific response”, a group of TFs with shared targets, each responding exclusively to a specific attack; “highly connected, multiple response” a group of TFs with highly connected targets, responding to different attacks (Figure 4B).

To identify those configurations, we used the ToMATo hierarchical clustering algorithm which builds the cluster hierarchy using a user-defined scalar-valued filter [37]. We defined two filters, one for the TF activities and one for the edge thickness, each following different mixtures of Gaussians. The process yields eight clusterings, one for each condition for the TF activity and one for the edges. The consensus clustering, obtained using the Cartesian product (see methods), yielded four main regions, corresponding to the four configurations (Figure 4C). We observe a tendency of TFs belonging to the same family to share targets, activity and response to pathogens (Supplementary Table 4).

Coupling the network structure and the activity estimation, our approach based on the Mapper and ToMATo algorithms from TDA yielded four configurations of how the tomato orchestrates the complex regulatory network to respond to different pests, identifying specific and common mechanisms.

### Four ERF and a NAC TFs coordinate the tomato response to multiple pathogens

To identify key regulators, we partitioned the global regulome accordingly to the four regions. Hence, we calculated degree and betweenness and the significance of the activity of TFs to retrieve the hubs of each region (see methods). We obtained five hubs, two in the green region and one in each of the other three; four belong to the ERF family and one to the NAC (Supplementary Table 5). These TFs were found to be active in response to pathogens with very different characteristics and/or infecting different tissues. The Solyc07g054220 and Solyc06g051840 in the green region actively respond to the virus severe strain, *B. cinerea* and *M. incognita* (14 dpi) and *C. fulvum*, respectively. Solyc03g093550 and Solyc12g009240 in yellow and red region, actively respond to *B. cinerea* and *M. incognita* (14 dpi) and *B. cinerea*, respectively. Finally, the Solyc07g63420 from the blue region, is activated upon *C. fulvum* and *B. cinerea* attacks. These findings may suggest that those five key regulators of plant defense are involved in a multi-pathogens response regardless pathogen characteristics or infected tissue. In support of that, a rising body of evidence shows that ERFs are closely involved in defense responses against various pathogens in plants [39] [40] [41].

To study if those hubs share targets, we extracted their first neighbors from the global GRN and we obtained a connected sub-network (Figure 5A). Targets can be shared by two or more hubs but none of the targets is targeted by the five TFs, and as expected the four ERFs share a very high number of targets (Figure 5B).

**Figure 5.**
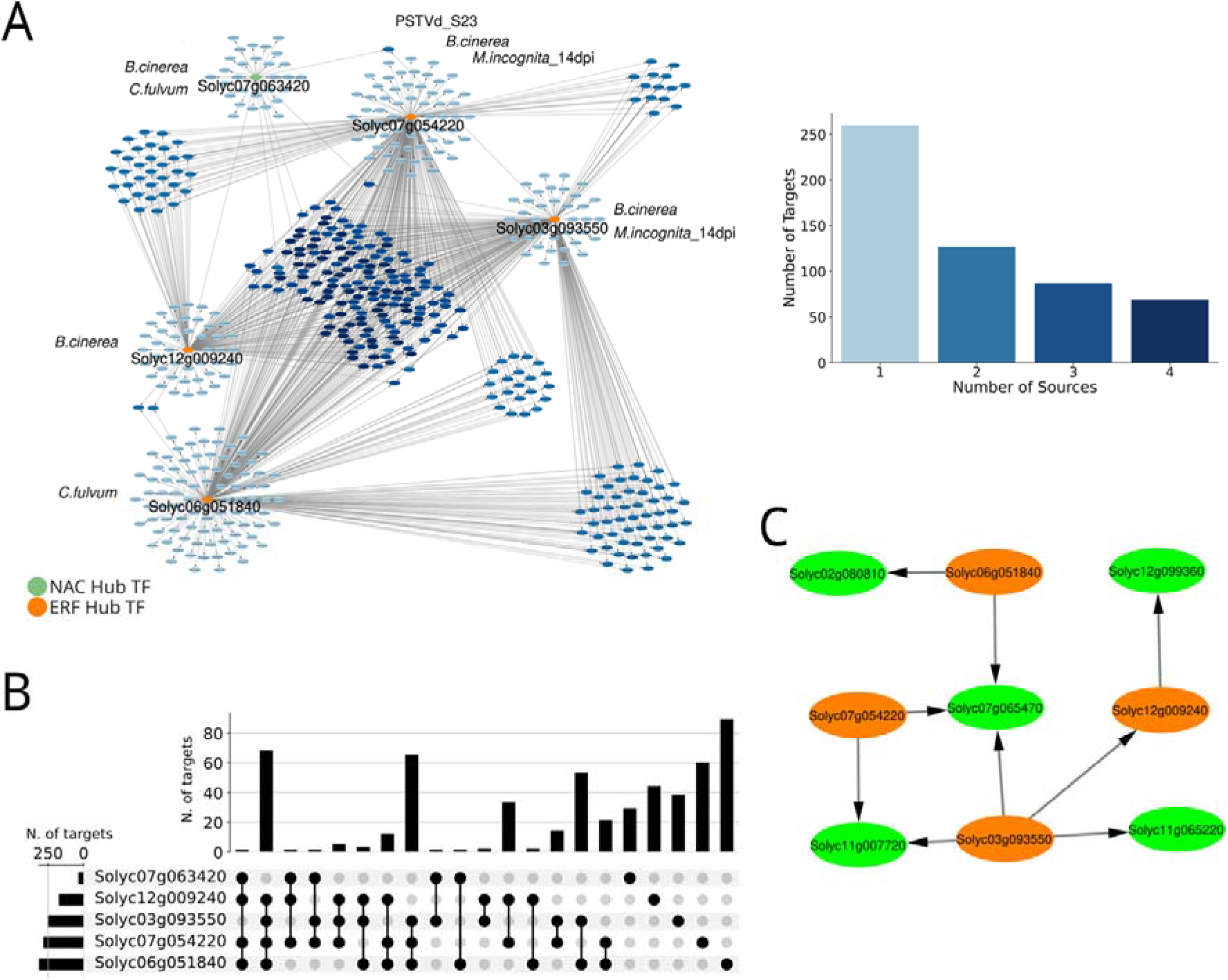
Hubs’ analysis on the four groups of clusters found by TDA. A) The sub network of the global GRN composed of the five hubs and their first targets. The conditions in which each TF was found with significant activity are represented next to the TF. The barplot represents the number of targets which are either specific to one TF or shared between multiple hubs from two to four. Colors of nodes in the network accordingly to the barplot. B) Upset plot showing the number of targets exclusive of each hub or shared among multiple hubs. On the left, barplot showing the total number of targets by hub. C) Sub-network of the TFs/targets associated to the pathway “carbon metabolism” with significant negative activity in response to *M. incognita*.

Since these TFs share several targets, we hypothesized that they may are involved in common biological functions. Therefore, we performed the pathway activity analysis (Supplementary Figure 7). Among the enriched pathways, the carbon metabolism captured our attention although the level of significance was close to the threshold, because of its pivotal role in plant response against biotic stress (Figure 5C). The pathway was found active in response to *M. incognita* aggression, including the four ERF hubs and five shared targets. Carbon metabolism and secondary metabolism are known to play a fundamental role in plants and tomato response to Meloidogyne species [42]. Furthermore, ERF TFs have been shown to be extensively associated with the regulation of secondary metabolites synthesis in response to biotic stresses [43] [44] [45]. Here, we found that ERF TFs may have a cooperative role in the modulation of this process during *M. incognita* infection. Furthermore, our study adds evidence of the role of this family in mediating the tomato response to multiple pathogens, expanding the putative role also in defense against *B. cinerea, C. fulvum* and PSTVd virus.

In conclusion, our analysis suggests that the four ERF hubs have a cooperative activity by sharing targets involved in pivotal pathways for plant defense and coordinate the tomato response to multiple bio-aggressors.

## Conclusions

Here we studied the response of the tomato to six distinct pathogens with a broad range of different characteristics from lifestyle to the infected tissue. The difficulty in realizing multi-stress experiments in laboratory, which increases with the number and the type of stress, has hindered the possibility to investigate plants’ molecular mechanisms in a more realistic scenario reproducing the field environment. To address this need, we developed novel *in silico* tools, to integratively analyze the complex GRN regulating plant molecular mechanisms to face multiple pathogen infections.

By jointly analyzing several transcriptomic data of tomato subjected to one pest at a time, we found regulatory mechanisms used to respond to either specific or multiple attacks. Although we observed that the general tomato response stratifies accordingly to infected tissues and subsequently with pathogen characteristics, by changing the scale of observation, we could highlight different mechanisms. We showed that TF activity is very complex, spanning from TFs with a significant activity triggered by multiple biotic agents, regardless of the pathogen’s features or infected tissue, to TF responding to only one attack. By creating a simplified representation of the original network, TDA algorithm allowed the identification of 18 clusters of TFs sharing targets either responding specifically to specific pests or having a cooperative or compensating activity by sharing targets and/or activity in response to multiple pathogens, or having an antagonist activity depending on the biotic stressor. By crossing the network structure and TF activity, TDA algorithms pinpointed four main regions in the GRN. Those regions permitted to shed light on how the tomato orchestrates the complex regulatory network to respond to different pests. The investigation of the properties of these four regions leads to the identification of five hubs belonging to the ERF family and one from the NAC family. Our findings suggest that those hubs play a key role in the coordination of regulatory mechanisms to face bio-aggressors. Overall, we demonstrated that TDA can provide innovative and robust instruments to analyze complex GRN.

Beyond the novel analysis methods proposed, we also developed another important resource for the plant scientists’ community: TomTom. With the wealth of data generated these days, it is extremely laborious to manually perform a thorough literature survey. TomTom yields a cartography of known interactions, all grouped together, from eleven databases. Overall, the interrogation of TomTom enables interactive exploration of current knowledge of molecular interactions in tomato and will help to provide mechanistic hypotheses and new insight into experimental observation. Furthermore, the workflow that we set up to build the knowledge graph is easily exportable for other plants; similarly, other interactions can be added for tomato in TomTom to expand the set of regulatory processes to be studied.

Our reproducible framework from knowledge graph to TDA can therefore be used on other biological models to study more generally host-pathogen interaction, but also in other fields of life sciences. In conclusion, our integrative and comparative approach enabled a more global understanding of tomato-pathogen interactions and will facilitate the development of multi-pathogen control strategies.

## Acknowledgments

This work was supported by the French government, through the UCA JEDI Investments in the Future project managed by the National Research Agency (ANR) under reference number ANR-15-IDEX-01. MC: This work was also partially supported by ANR grant “TopModel”, ANR-23-CE23-0014, and supported by the French government, through the 3IA Cote d’Azur Investments in the project managed by the National Research Agency (ANR) with the reference number ANR-23-IACL-0001. This project was supported by INRAE in the framework of the “Métaprogramme DIGIT-BIO” and Plant Health and Environment department.

We are grateful to the bioinformatics and genomics platform, BIG, Sophia Antipolis (ISC PlantBIOs, https://doi.org/10.15454/qyey-ar89) for computing and storage resources, and we would like to thank Martine Da-Rocha for her meaningful guidance regarding the RNA-seq pre-processing.

## Authors’ contributions

MM: methodology; software; validation; formal analysis; resources; data curation; writing - original draft; visualization. MC: formal analysis. XAG: formal analysis; resources. ADa: software. SL: methodology; software. JSR: review & editing; supervision. SJP: conceptualization; writing - review & editing. ADu: methodology; writing - review & editing; supervision. SB: conceptualization; methodology; writing - original draft; supervision; project administrator; funding acquisition.

